# Regions in the human inferior temporal gyrus are engaged in numerosity processing across visual stimulus categories

**DOI:** 10.1101/2025.08.21.671518

**Authors:** Vicente Alejandro Aguilera González, Gizem Cetin, Stephanie Zika, Antonia Schulz, Max Pirsch, Alexia Dalski, Mareike Grotheer

## Abstract

The visual number form areas (here referred to as ITG-math) have gained recent attention as a neural substrate of mathematical cognition. Yet, the function of these regions remains debated with some research suggesting selectivity for digits and others suggesting a visual stimulus-independent role in mathematical processing. Here we addressed this debate by exploring the impact of different tasks and stimuli on neural responses in ITG-math. Participants were presented with digits and UFOs, while performing a 1-back task either on stimulus numerosity or on stimulus color. We collected two sessions of this experiment from 17 healthy adults and could identify voxels with higher responses in the numerosity than the color task in 92% of the participants, whereas contrasting digits with UFOs lead to selective responses in only 35% of the participants. Accordingly, in independent data, ITG-math showed a task preference, but no stimulus preference. Multivariate analyses further revealed task encoding, and a combination of task and stimulus encoding, in the left and right ITG-math, respectively. Our work hence supports the notion that the role of ITG-math in mathematical cognition goes beyond the encoding of digits.

## 1. Introduction

Mathematical abilities are critical in our daily lives, yet our current understanding of the neural substrates of these abilities is limited. Bilateral functional regions located in the posterior inferior temporal gyrus (pITG) have gained attention recently for their preference to visually presented digits (i.e. Arabic Numerals) over other visual stimuli (Pinel et al. 1999; Roux et al. 2008; Cui et al. 2013; Shum et al. 2013; Hermes et al. 2015; Amalric and Dehaene 2016; Daitch et al. 2016; M. Grotheer et al. 2016). Applying transcranial magnetic stimulation to these regions causes a significant impairment in the identification of digits (Mareike Grotheer et al. 2016), suggesting a causal role in digit processing. Due to their preference for digits over other stimulus categories, these functional regions are often referred to as the Visual Number Form Areas (VNFA) or ITG-numbers.

Importantly, the functional role of ITG-numbers remains debated. While several studies have successfully identified a category-selective region for digits in the pITG (Pinel et al. 1999; Roux et al. 2008; Cui et al. 2013; Shum et al. 2013; Abboud et al. 2015; Mareike Grotheer et al. 2016; M. Grotheer et al. 2016), various others found no category-selectivity (Pinel et al. 2004; Libertus et al. 2009; Cappelletti et al. 2010; Andres et al. 2012; Attout et al. 2014; Carreiras et al. 2015; Cummine et al. 2015; Holloway et al. 2015; Peters et al. 2015; Merkley et al. 2019). Moreover, there is mounting evidence that math-related tasks induce increased neural responses in this part of the cortex, irrespective of the visual features and even the modality in which these tasks are presented (Abboud et al. 2015; Hermes et al. 2015; Amalric and Dehaene 2016; Grotheer et al. 2018), suggesting that regions in the pITG may be involved in mathematical cognition beyond the processing of visual digits. As a middle ground between these views, it has also been suggested that the pITG might contain two distinct functional subregions (Daitch et al. 2016): a smaller and more anterior region selective for digits (ITG-numbers), and a larger and more posterior region that is selectively involved in math tasks irrespective of the visual stimulus these tasks are performed on (ITG-math). Overall, the functional role of regions in the pITG remains debated and further research is needed to disentangle the impact of visual stimulus features and mathematical task demands on their neural responses.

In the current study we aimed to address this debate by manipulating both the visual stimulus features and the task demands orthogonally within the same fMRI paradigm. Participants (N=17) were presented with numerosities between 1 and 5 either as digits or as UFOs while they performed either a numerosity or color (control) task (Fig. 1) in two experimental sessions.

**Fig. 1.**
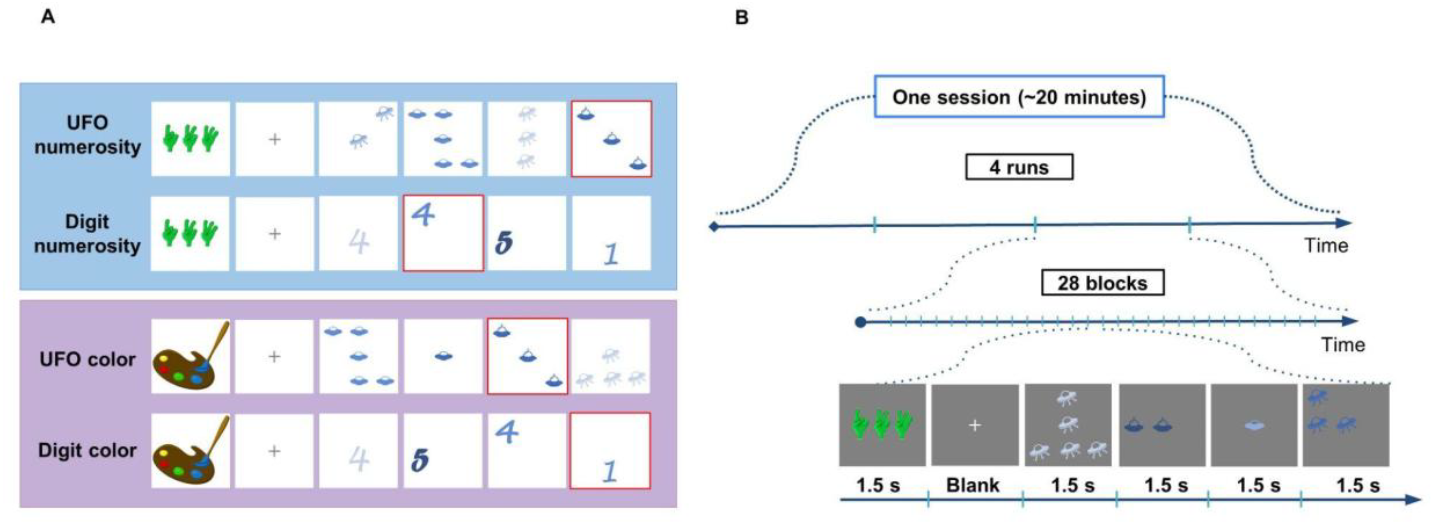
Experimental design. A. Overview of the experimental conditions. Subjects viewed UFOs or digits (Arabic numerals) on a gray background while performing a 1-back task attending either to the numerosity or the color of the stimuli. B. Overview of the block structure. The experiment comprised four runs, each containing 28 blocks. At the beginning of each block, a cue indicating the task was presented for 1.5 seconds. This was followed by a blank period of 0, 1.5 or 3 seconds. Immediately after, four images of a single experimental condition were presented for 1.5 seconds each.

Our analyses revealed 4 main findings: 1) functional regions of interest could be identified reliably within individuals when contrasting the numerosity with the color task (ITG-math) but not when contrasting digits with UFOs (ITG-numbers), 2) the ITG-math subregions showed a preference for the numerosity task in independent data, 3) the spatial location of ITG-math was consistent across session, and 4) distributed responses in ITG-math suggest task encoding in the left hemisphere, and task and stimulus encoding in the right hemisphere. Overall our results hence suggest that regions in the pITG are involved in numerosity processing independently of the visual stimulus features of the presented stimuli. Moreover, we propose that the developed experimental paradigm may serve as a localizer to identify numerosity-selective responses in the pITG and to hence facilitate further research on these regions.

## 2. Materials and Methods

### 2.1. Participants

Nineteen adult volunteers (12 females, mean age 27.5 ± 4,16 years, 17 right-handed) participated in two separate sessions on different days. Before scanning, participants were instructed on the task they were expected to perform and received a brief training on a laptop. Two participants chose not to continue the study after their first session. No participants were excluded due to participant motion (head motion between or within runs above 3 voxels in any direction). The final data set was hence composed of 17 participants (11 females, mean age 27 ± 4,15 years, 15 right-handed). Subjects had normal or corrected to normal vision and gave their informed written consent. The procedure was approved (approval ID: AZ 159/21) by the Philipps-University Marburg’s Ethics Committee from the Medicine Faculty.

### 2.2. Data acquisition and preprocessing

Acquisition: Data was collected using a Siemens 3 tesla Magnetom Prisma scanner with a 64-channel head coil. We acquired 42 slices covering the whole brain using two T2*-sensitive gradient echo-planar sequences. One for the main experiment (voxel size: 2.5 mm x 2.5 mm x 2.5 mm, TR: 1500 ms, TE: 30 ms, FoV: 192 mm, flip angle: 71°, slice thickness: 2.5 mm, acceleration factor of 2) and another for the localizer (voxel size: 2.5 mm x 2.5 mm x 2.5 mm, TR: 1000 ms, TE: 30 ms, FoV: 192 mm, flip angle: 59°, slice thickness: 2.5 mm, acceleration factor of 3). Additionally, a whole-brain, anatomical volume was collected once for each participant using a T1-weighted BRAVO pulse sequence (Voxel size: 0.9 mm x 0.9 mm x 0.9 mm, TI: 949 ms, FoV: 240 mm, flip angle: 8°, slice thickness: 0.94 mm).

Processing: The anatomical brain volume of each subject was segmented into gray and white matter using FreeSurfer 7.3.2 (https://surfer.nmr.mgh.harvard.edu/), with manual corrections using ITKGray (https://github.com/vistalab/itkgray), and each participant’s cortical surface was reconstructed. Functional data was analyzed using the mrVista toolbox (https://github.com/vistalab) for Matlab (R2021b). fMRI data from each experiment was motion-corrected within and between runs, and then aligned to the T1-weighted anatomical volume. We did not apply any spatial smoothing. In each experiment, a separate design matrix was created and convolved with a hemodynamic response function in Matlab to generate predictors for each experimental condition. We estimated regularized response coefficients (betas) for each voxel and each predictor using a general linear model (GLM) indicating the magnitude of response for that condition.

### 2.3. Experimental design

#### Main experiment (UFO-paradigm)

##### Stimuli and tasks

We presented images of digits or UFOs during either a numerosity or a color task. Stimulus numerosities ranged from 1 to 5, and stimuli were depicted in one of five possible hues of blue. Behavioral pilots were used to select color hues in such a way that task difficulties in the numerosity and color task were matched as closely as possible. Three different types of UFOs and three different fonts for the digit images were used, resulting in a total of 75 unique images per stimulus category. Participants performed two different 1-back tasks on the same stimuli: Numerosity: Participants were instructed to press a button whenever two consecutive images within a block depicted the same numerosity. Color: Participants had to press a button whenever two consecutive images repeated the same hue of blue. A cue presented prior to each block indicated the participant’s task. For the numerosity task, the cue image displayed three alien hands indicating different numerosities, whereas a painter’s palette was used for the color task (Fig. 1A).

##### Trial structure

The experiment was a mixed design with four experimental conditions: UFO-numerosity, digit-numerosity, UFO-color, and digit-color. Four ∼5 min long runs were collected in each session with each run containing 28 blocks from a single condition (Fig. 1B). Cues lasted 1500 ms and were followed by a blank period of either 0, 1500 or 3000 ms. In each block, four images were presented for 1500 ms each, with an inter-stimulus interval (ISI) of 0s. To isolate and remove activations induced by the cues, four “blank blocks” were included in which only the cues were presented. All stimuli covered a visual angle of ∼2.5°. The experimental conditions were pseudo-randomized so that each condition appeared six times per run.

##### Color Localizer

Color-selective control regions in the ventrotemporal cortex (VTC) were identified via a visual localizer experiment designed in previous work (Stigliani et al. 2015; Grotheer et al. 2018; Dalski et al. 2024). In this localizer, five visual stimulus categories were presented either in color or in grayscale: faces, houses, limbs, text, and objects. Each stimulus was presented for 500 ms, with an ISI of 0 s, in blocks of 8 stimuli. Each participant completed 4 runs that were ∼5 min long and included 10 repetitions of each stimulus category. All stimuli subtended a visual angle of ∼8°. The task was to fixate the center of the screen and to press a button when only the background without any image was presented (oddball task). 3% of all stimuli were oddballs.

### 2.4. Regions-of-interest definition

Functional regions of interest (fROIs) were defined in the native brain space of each participant, using both functional and anatomical criteria. The first session of the main experiment and the color localizer data were used to define the fROIs. These were then used to evaluate responses in the second session ensuring independence between fROI definition and data analysis.

#### ITG-math and ITG-numbers

The current literature indicates a preference for number stimuli(Roux et al. 2008; Shum et al. 2013; Mareike Grotheer et al. 2016) and mathematical processing (Hermes et al. 2015; Amalric and Dehaene 2016; Daitch et al. 2016; Grotheer et al. 2018) in the pITG. Thus, we aimed to define fROIs for both selectivities using data from the main experiment (T=3, voxel level, uncorrected). First, we contrasted the digit with the UFO stimuli, to identify regions selective for digit stimuli (ITG-numbers). ITG-numbers (average size: left hemisphere: 198.86 (±22.24) mm3, right hemisphere: 124.40 (±32.27) mm3) was found in 7 (41.18%) participants in the left and 5 (29.41%) participants in the right hemisphere. As ITG-numbers could not be reliably identified from contrasting digit stimuli with UFO stimuli using all data, we also tried to define the regions contrasting digit stimuli with UFO stimuli only within the numerosity task. In this case, ITG-numbers (average size: left hemisphere: 106.80 (±33.31) mm3, right hemisphere: 43.50 (±16.50) mm3) was found in 6 (35.30%) participants in the left and 2 (11.76%) participants in the right hemisphere.

Next, we contrasted responses to the numerosity with the color task, to identify regions selective for numerosity processing (ITG-math). ITG-math was identified in 16 participants in the left and 15 participants in the right hemisphere. On average, the size of left and right ITG-math was 303 (±48.89) mm3 and 380 (±73.25) mm3, respectively. As ITG-math regions were more reliably identifiable than ITG-numbers regions, we focused on ITG-math for most analyses. Additionally, a second ITG-math was defined using responses to the numerosity task in the second session, in order to calculate the spatial consistency across sessions (average size: left hemisphere: 434.93 (±93.30) mm3, right hemisphere: 432.23 (±107.20) mm3). To test the efficiency of the developed paradigm in identifying ITG-math and hence its applicability as a localizer, we repeated the same procedure using data from only the first two runs in each session (i.e. ∼10 minutes worth of data). With this shorter experimental paradigm, ITG-math in session 1 (average size: left hemisphere: 231.69 (±52.05) mm3, right hemisphere: 230.28 (±49.60) mm3) could be defined in 13 (76.47%) and 14 (82.35%) participants in the left and right hemisphere, respectively. In session 2 ITG-math (average size: left hemisphere: 182.15 (±31.12) mm3, right hemisphere: 183.07 (±42.00) mm3) could be defined also in 13 (76.47%) and 14 (82.35%) participants in the left and right hemisphere, respectively.

#### Color-sensitive region

We contrasted responses to colored and gray-scale images in the color localizer data to define three color-sensitive patches in the medial fusiform gyrus (Lafer-Sousa et al. 2016). We identified the anterior (Ac-color), central (Cc-color), and posterior color regions (Pc-color) in 15, 17, and 17 participants in the left hemisphere, and in 16, 17, and 16 participants, respectively in the right hemisphere (average size Ac-color: left hemisphere: 414.67 (±61.87) mm^3^, right hemisphere: 457.37 (±90.45) mm^3^; Cc-color: left hemisphere: 755.23 (±149.37) mm^3^, right hemisphere: 754.23 (±116.82) mm^3^; and Pc-color: left hemisphere: 753.94 (±141.84) mm^3^, right hemisphere: 936.62 (±156.86) mm^3^).

### 2.5. Statistical analysis

#### Behavioral responses

We assessed subjects’ performance (measured in % correct) as well as reaction times (RTs, measured in ms) with repeated measures ANOVAs (rmANOVAs). In the main experiment we used task, stimulus, and session as factors. In the color localizer, we used stimulus and color as factors. Tukey-Kramer post-hoc tests were applied when the rmANOVAs revealed significant main effects or interaction effects.

#### Univariate analysis

For both the main experiment and the color localizer, the average time course in percentage signal change (PSC) was extracted for each condition from each fROI. Then we applied GLMs to estimate betas (measured in units of PSC±SEM; Fig. 2; Supplementary Fig. 1), which indicate the magnitude of the response for each condition. Crucially, we used independent data for fROI definition (main experiment session 1 for ITG-math, color localizer for color regions) and signal extraction (main experiment session 2). We analysed univariate responses in the main experiment with rmANOVAs that used hemisphere, task, stimulus, and session as factors. These analyses were also replicated using the 2-run ITG-math fROI and independent data from session 2 (Supplementary Fig. 2).

**Fig. 2.**
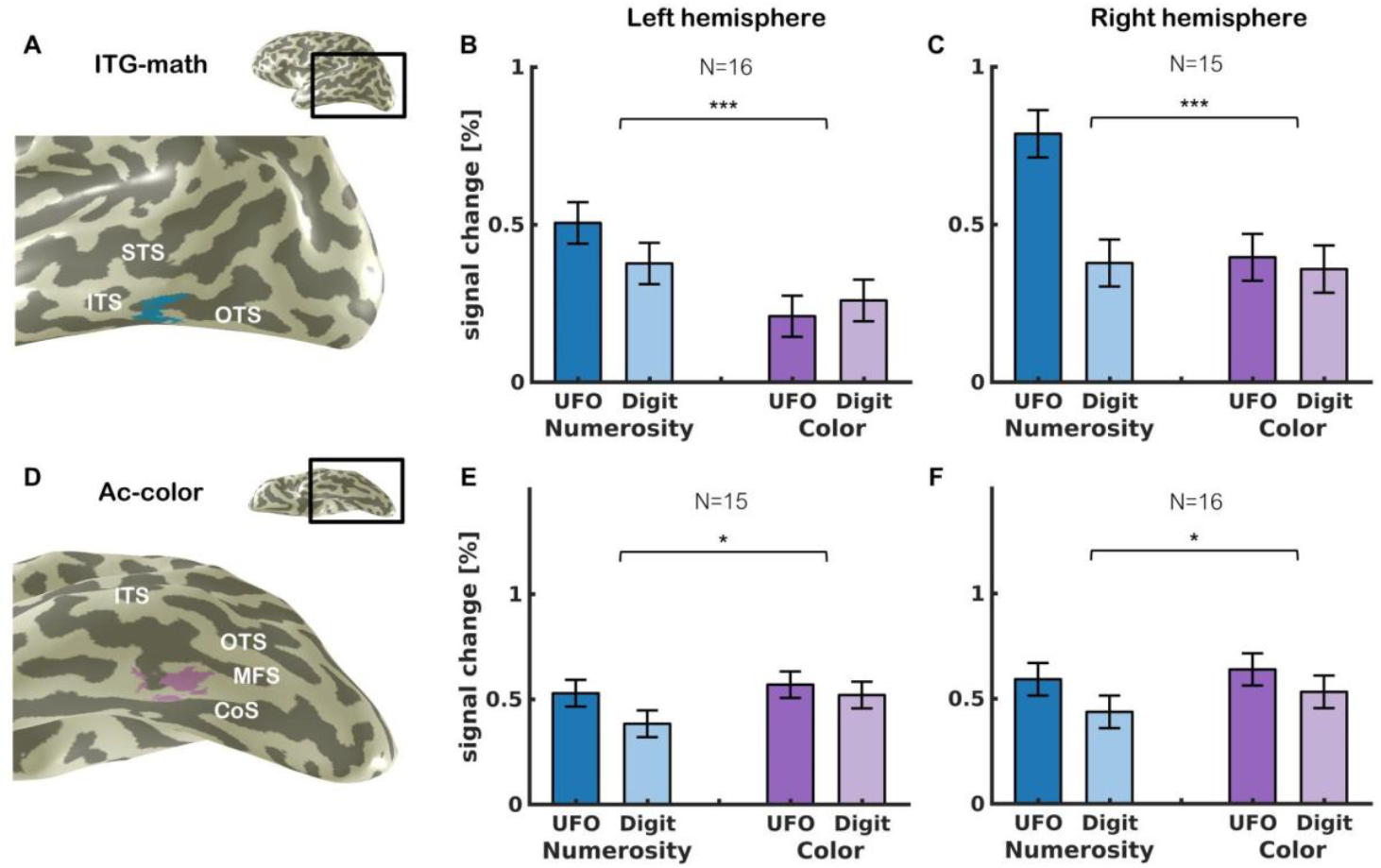
ITG-math shows a preference for the numerosity task. A. ITG-math: voxels in the ITG/ITS that showed significantly higher responses during the numerosity than the color task (T=3, voxel level, uncorrected). Data is shown in a lateral view of the inflated left hemispheres of a representative subject. B. Mean responses ± SEM across 16 subjects in the left ITG-math from independent data. C. Mean responses ± SEM across 15 subjects in the right ITG-math from independent data. D. Ac-color: voxels in the median fusiform gyrus (FG) that showed significantly higher responses for colored than gray-scale images in the color localizer (T=3, voxel level, uncorrected). Data is shown in a ventral view of the inflated left hemispheres of a representative subject. E. Mean responses ± SEM across 15 subjects in the left Ac-color. F. Mean responses ± SEM across 16 subjects from the right Ac-color. Asterisks indicate main effect of task across hemispheres, *p<0.05, ***p<0.001. Abbreviations: STS: superior temporal sulcus, ITS: inferior temporal sulcus, OTS: occipitotemporal sulcus, MFS: midfusiform sulcus, CoS: collateral sulcus.

#### Dice coefficient (DC)

To evaluate the spatial reliability of ITG-math across sessions, we first defined the anatomical boundaries of the posterior ITG, i.e. the area of cortex in which ITG-match could fall: anteriorly: the posterior hippocampus; inferiorly: the occipito-temporal sulcus (OTS); superiorly: the middle temporal gyrus (MTG); and posteriorly: the anterior occipital sulcus (AOS)). We then evaluated the spatial overlap of ITG-math defined in the two sessions within these boundaries and quantified it with the Dice coefficient 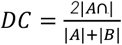, where |A| and |B| are ITG-math from session 1 and session 2 data, respectively, and |A ∩ B| is the intersection between them. The DC quantifies the similarity of two samples from 0 to 1, indicating none or a complete overlap, respectively (Dice 1945). To assess if the measured DC was above chance, we generated a chance level by randomly positioned 2 disk ROIs, matched in size to ITG-math from session 1 and 2, within a square matching the size of the combined anatomical ITG and quantified the DC. This process was repeated 10000 times within each subject, and the average DC was defined as the chance level DC. We then used paired t-tests to evaluate if the measured DC was different from chance (Fig. 3B). The same procedure was replicated for ITG-math fROIs defined from two runs in session 1 and 2 (Supplementary Fig. 3).

**Fig. 3.**
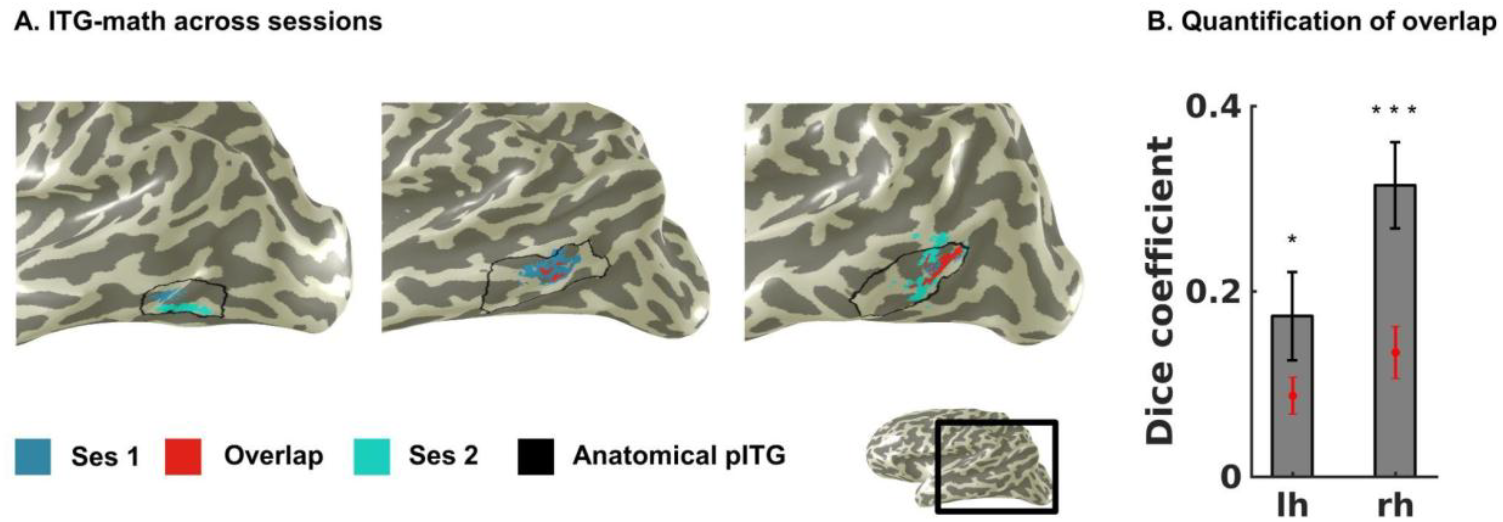
ITG-math fROIs show consistent localization across sessions. A. Figure shows the inflated cortical surface of three participants representing examples of low (left), medial (middle), and high (right) spatial overlap in ITG-math fROIs across sessions. B. Quantification of the overlap between fROIs across sessions using the dice coefficient (DC), mean across subjects ± SEM. Red dots indicate chance level. Star indicates DCs is significantly higher than chance, *p<0.05, ***p<0.001.

#### Multivoxel pattern analysis

Disk fROIs, centered on their respective fROI, but with constant radii that matched the average fROI sizes across participants (ITG-math: left hemisphere: 7.14 mm; right hemisphere: 7.80 mm; right Ac-color: 8.10 mm) were used for the multivoxel pattern analyses (MVPAs, Fig. 4). First, the responses to each experimental condition were computed for each voxel with a GLM. Responses were then normalized and z-transformed. Subsequently, we used a leave-one-run-out procedure to calculate correlations between pairs of multivoxel patterns (MVPs), and summarized them in representation similarity matrices (RSMs), which were first calculated at subject level and then averaged across subjects (Fig. 4D, E). Next, we compared the measured group-level RSMs to model RSMs for task, stimulus, and combined task using Spearman’s correlation as implemented in the MNE RSA toolbox (Vliet et al. 2025: https://github.com/mne-tools/mne-rsa). Winner-takes-all (WTA) classifiers, with a leave-one-run-out procedure, determined in each subject and experiment, whether stimulus and/or task information can be decoded from MVPs in the fROIs. Paired t-tests were used to evaluate if WTA classifier performance exceeded 50% chance level (Supplementary Fig. 4). The same procedure were replicated using the native functionally defined ROIs of each individual (Supplementary Fig. 5) as well as with disk fROIs that were defined from the 2-run fROIs following (average 2-run fROI size: left hemisphere: 6.20 mm; right hemisphere: 6.40 mm, Supplementary Fig. 6).

**Fig. 4.**
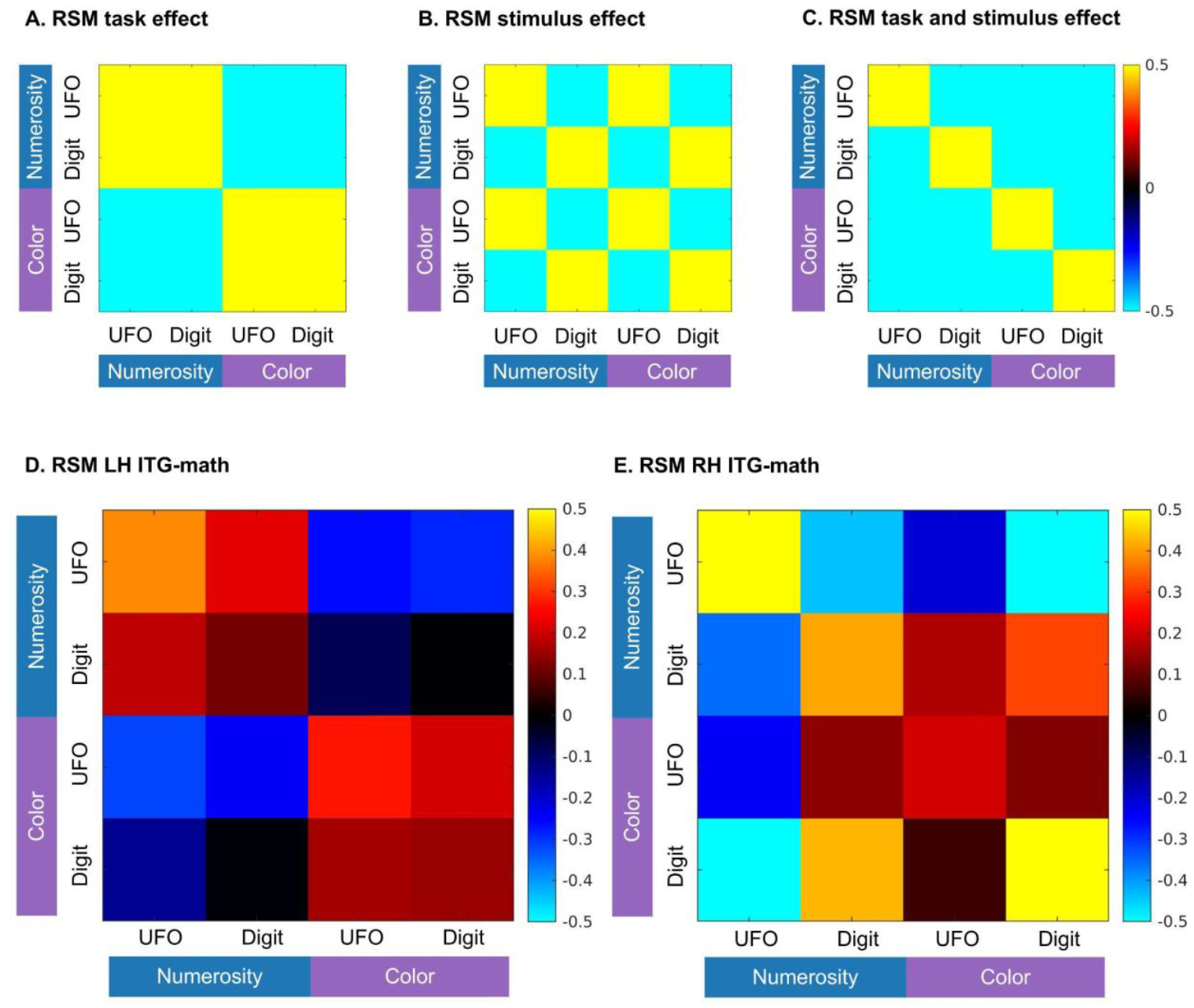
Distributed responses in ITG-math vary across hemispheres. A-C. Model RSMs depicting hypothetical similarities across conditions based on task (A), stimulus (B), and combined task and stimulus effects (C). D-E. RSM from ITG-math in the left (D. N=16) and right (E. N=15) hemispheres. Conditions are arranged by stimulus (UFO vs. digit) and grouped by task (numerosity vs.color).

#### Group-level whole brain analysis

We also evaluated whole brain responses in the main experiment by combining the contrast maps of interest (numerosity vs color task, and digit vs UFO stimuli) from both sessions at subject-level. The combined contrast maps of all subjects were mapped to the FreeSurfer average brain (T≥1 at voxel level, uncorrected) for visual inspection (Fig. 5, Supplementary Fig. 7). The same procedure was repeated with contrast maps created using only the first two runs of session 1 (Supplementary Fig. 8).

**Fig. 5.**
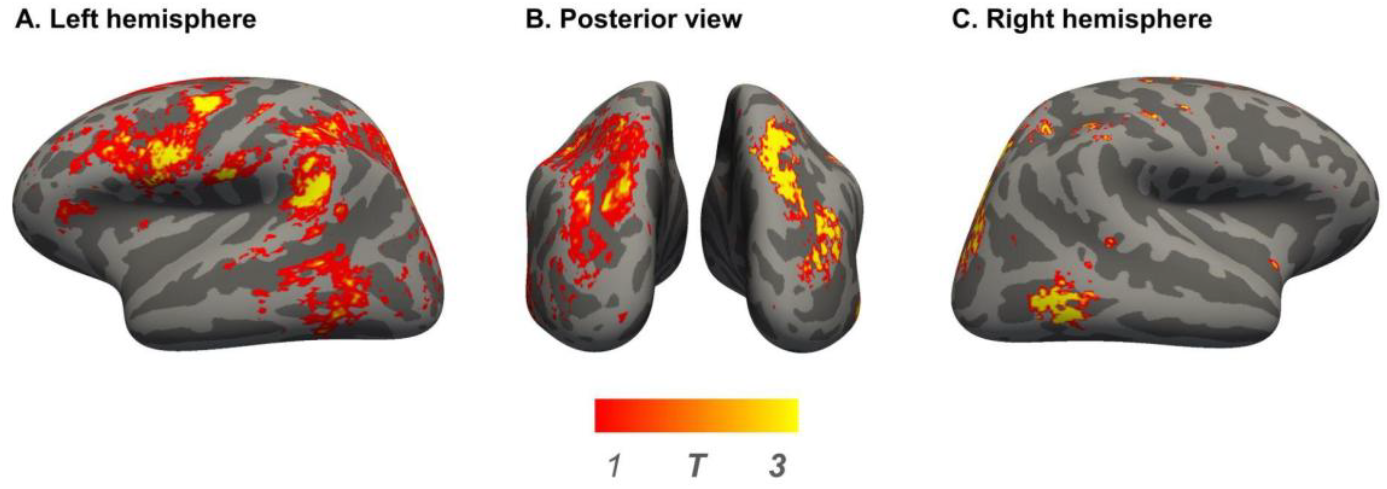
Whole-brain analysis contrasting the numerosity and color tasks across participants. Responses elicited by contrasting the numerosity with the color task in both sessions were averaged across participants and are presented on the inflated cortical surface of the Freesurfer average brain.

#### Code and data availability

Experimental and analysis code can be found in GitHub: https://github.com/EduNeuroLab/ITG-math_localizer.git MRI data will be made available by the corresponding author upon reasonable request.

## 3. Results

### 3.1. All tasks were performed with high accuracy

Here we tested whether a numerosity task and/or digit stimuli can be used to reliably identify functional subregions in the pITG that are involved in mathematical processing. Seventeen healthy adults were presented with UFOs and digits, while they performed either a numerosity or a color task. We employed rmANOVAs to compare performance and reaction times (RTs) across conditions and sessions. In performance, these rmANOVAs revealed a task effect (F(1,16)=4.76, p=0.04) with higher performance in the numerosity task 95.23% (±0.02%) than the color task 91.73% (±0.02%). There was no significant stimulus (F(1,16)=1.45, p=0.24) or session effect (F(1,16)=0.53, p=0.47), but an interaction effect of task and stimulus was found (F(1,16)=6.56, p=0.02), with highest accuracies for digits in the numerosity task (97.55±2.11%). In the RTs, rmANOVAs revealed significantly shorter RTs in the color task than the numerosity task (F(1,16)=14.10, p=0.002; color: 702 (±27) ms; numerosity: 760 (±26) ms). There was no significant stimulus (F(1,16)=1.06, p=0.32) or session effect (F(1,16)=2.79, p=0.11). An interaction effect of task and stimulus (F(1,16)=13.21, p=0.002) was found for RTs; in the numerosity task, RTs were shorter for numbers (730 (±26) ms) than UFOs (790 (±26) ms) (p=0.03), whereas the opposite was found for the color task (p=0.03, numbers: 717 (±28) ms, UFOs: 688 (±27) ms).

In addition to the main experiment, we also collected a color localizer to identify color-sensitive regions in the medial fusiform gyrus that were used as control ROIs. In this localizer, rmANOVAs revealed no effects of color or stimulus and no interaction effect between color and stimulus in either performance (color: F(1,16)=3.21, p=0.09; stimulus: F(4,64)=2.08, p=0.09, interaction: F(4,64)=0.54, p=0.71) or RTs (color: F(1,16)=2.13, p=0.16; stimulus: F(4,64)=0.74, p=0.57, interaction: F(4,64)=0.74, p=0.57).

### 3.2. ITG-math shows stronger activations in the numerosity than the color task

We started by contrasting the numerosity and color tasks, and the digit and UFO stimuli using only session 1 data (T≥3 voxel-level, uncorrected). When contrasting tasks, 16 and 15 of the subjects (total N=17) showed selective activations for the numerosity task in the left and right posterior ITG, respectively (for example subject see Fig. 2A, B). In contrast, when comparing responses to different stimuli across tasks, only 7 and 5 of the subjects showed a preference for digits in the left and right posterior ITG, respectively. We also tested for a preference for digits only within the numerosity task, however in this analysis only 6 and 2 of the subjects showed a preference for digits in the left and right pITG, respectively.

As reliable activations across subjects were only found for the task contrast, we focus on the fROIs defined by their task preference in the following analyses and we refer to these fROIs as ITG-math. Across participants, ITG-math was located posterior to the fMRI signal dropout zone near the ear canal (Supplementary Fig. 9, 10). Within each participant, we extracted ITG-math’s responses in the second session (i.e. independent data) and assessed its response profile using rmANOVAs with task, stimulus, and hemisphere as factors. We found a main effect of task (F(1,13)=33.30, p<0.0001), with higher responses in the numerosity task, as well as a main effect of stimulus (F(1,13)=13.33, p=0.003), with higher responses for the UFO stimuli. We also found an interaction effect between hemisphere and stimulus (F(1,13)=13.11, p=0.003). Subsequent post-hoc tests revealed significantly higher responses to UFOs than digits in the right hemisphere (p=0.0002; Fig. 2B) but not the left hemisphere (p=0.50). No other main effects or interaction effects were found. To evaluate the efficiency of the experimental paradigm in localizing ITG-math we reproduced these analyses with ITG-math fROI defined from only the first two runs of session 1 (i.e. 10 minutes worth of data) and similarly found a main effect of task (F(1,11)=21.48, p=0.0007), a main effect of stimulus (F(1,11)=24.24, p=0.0004) (Supplementary Fig. 2).

In our main control region Ac-color, we found a main effect of task (F(1,14)=4.99, p=0.04) with higher responses in the color task, as well as a main effect of stimulus (F(1,14)=12.15, p=0.004), with higher responses for UFO stimuli. No main effect of hemisphere or interaction effects were found. For results of additional control regions Cc-color and Pc-color please see Supplementary Figure 1.

### 3.3. ITG-math shows a consistent spatial location within individuals across sessions

To determine if ITG-math’s spatial location is consistent across sessions, next, we also defined ITG-math by contrasting the numerosity and color tasks in session 2 data (T≥3 voxel-level, uncorrected). We could define an ITG-math fROI in most of the participants (left hemisphere: N=14; right hemisphere: N=13). Across sessions, ITG-math fROIs overlapped in 12 of the subjects in both hemispheres. The average DCs (mean across subjects ± standard error (SE): left hemisphere: 0.17 (±0.05), right hemisphere: 0.31 (±0.05)) were significantly above chance in both hemispheres (left: p=0.02; right: p=0.0007; Fig. 3B). An additional independent t-test across hemispheres revealed significantly higher DCs in the right hemisphere (p=0.04). Significantly above chance DCs in both hemispheres (left: p<0.0001; right: p=0.05) were also obtained when we only used data from the first two runs to define ITG-math in each session (Supplementary Fig. 3).

### 3.4. Distributed responses in ITG-math differ across hemispheres

The univariate analyses described above revealed that regions in the pITG are selectively engaged in numerosity processing, but not in processing digits over other visual stimuli that convey numerical information. Nonetheless, it is still plausible that information about visual stimulus features is represented on a finer spatial-scale and extractable from differences in the response patterns across voxels within the pITG regions. To test this possibility and to determine what kind of information is encoded within the pITG-regions, we also conducted multi-voxel pattern analyses (MVPAs) in session 2 data using constant size disk ROIs centered on session 1 ITG-math fROIs. We computed representational similarity matrices (RSMs) for each subject using a leave-one-run out procedure and then compared the group-averaged RSMs of each hemisphere to three model RSMs: 1) task, 2) stimulus, and 3) combined task and stimulus (Fig. 4A, B, C). In the left ITG-math, we found a significant correlation only with the task-model RSM (task: r=0.87, p<0.0001; stimulus: r=0.03, p=0.92; task and stimulus r=0.50, p=0.05). In the right ITG-math, a significant correlation with the stimulus model, and with the combined task and stimulus model was found (stimulus: r=0.65, p=0.006; task and stimulus: r=0.66, p=0.006), but no significant correlation with the task model was observed (r=0.24, p=0.36). Winner-takes-all (WTA) classifiers showed similar result patterns across the two hemispheres (Supplementary Fig. 4A, B). RSAs in native fROIs and constant-disk fROIs defined based on only the first 2 runs of session 1 showed similar results to the constant size disk ROIs with significant correlation with the task model in the left hemisphere (native fROI: r=0.87, p<0.0001, Supplementary Fig. 5D; 2-run disk fROI: r=0.87, p<0.0001, Supplementary Fig. 6D), and with the stimulus model in the right hemisphere (native fROI: r=0.54, p=0.03, Supplementary Fig. 5E; 2-run disk fROI: r=0.54, p=0.03, Supplementary Fig. 6E). Additionally, native fROIs in the right hemisphere also significantly correlated with the task and stimulus model (r=0.63, p=0.009, Supplementary Fig. 5E). WTA classifiers trained on MVPAs from native and 2-run disk fROIs yielded similar results (Supplementary Fig. 5F, G; Supplementary Fig. 6F, G, respectively).

### 3.5. ITG-math is part of a network of brain regions preferentially engaged in numerosity processing

Here we found a reliable preference for the numerosity task in the posterior ITG, which raises the question if other regions of the brain may also show such a preference. To test this, we averaged individual subjects’ contrast maps comparing the numerosity and color tasks and evaluated the resulting group-average activation maps. As expected, visual inspection of the group maps revealed a preference for the numerosity task in the pITG bilaterally. In addition, in the left hemisphere, a preference for the numerosity task was observed in the intraparietal sulcus (IPS), the supramarginal gyrus (SMG), and the precentral sulcus (PCS) (Fig. 5A, B), while in the right hemisphere, the preference for the numerosity task was largely limited to the pITG and the IPS (Fig. 5B, C). Averaged contrast maps (numerosity vs color) from only the first two runs of session 1 showed a numerosity preferences in the same regions (Supplementary Fig. 8). A group average map contrasting responses to digits and UFOs is presented in Supplementary Fig. 7.

## 4. Discussion

In the current study, we addressed an ongoing debate about the functional role of regions in the posterior ITG that have been implicated in mathematical cognition: Are these regions selectively involved in processing digits or do they play a broader - stimulus independent - role? To test this, we manipulated participants’ tasks (numerosity task vs color task) and the visual stimulus tasks were performed on (digits vs UFOs) orthogonally within the same experiment. Our analyses revealed 4 main findings: 1) functional regions of interest could be identified reliably within individuals when contrasting the numerosity with the color task (ITG-math) but not when contrasting digits with UFOs 2) the ITG-math regions showed a preference for the numerosity task in independent data, 3) the spatial location of ITG-math was consistent across session, and 4) distributed responses in ITG-math revealed task encoding in the left hemisphere, and task and stimulus encoding in the right hemisphere.

We could successfully identify ITG-math in 92% of the participants by contrasting responses in the numerosity and color tasks. These results are consistent with previous studies finding a format-invariant selectivity for mathematical tasks in the pITG (Coderre et al. 2009; Cui et al. 2013; Grotheer et al. 2018; Cai et al. 2021; Cai et al. 2023). Importantly, ITG-math displayed reproducible localization, as measured with DCs, which is also considered an indicator of specialisation (Cohen and Dehaene 2004). Interestingly, responses to UFOs were higher than to digits across hemispheres during the numerosity task. We hypothesize that these differences could be driven by the more resource-demanding nature of non-symbolic numerosities (Skagenholt et al. 2018; Sokolowski et al. 2022), which is consistent with the higher RTs observed for UFOs in the numerosity task. Altogether, these findings indicate that the pITG contains a functional subregion engaged in numerosity tasks (ITG-math), which may support mathematical cognition irrespective of the stimulus. As indicated by our whole-brain analyses, ITG-math is part of a larger network of brain regions involved in numerosity processing which includes the bilateral IPS, and left SMG and PCS, i.e. regions which have consistently been associated with mathematical cognition in prior work (Menon et al. 2000; Daitch et al. 2016; Damarla et al. 2016; Nakai and Nishimoto 2023). We were able to identify ITG-math and the broader numerosity network even when using only 2 runs (i.e. 10 minutes worth) of data suggesting that the experimental paradigm developed here, which is publicly available, may serve as an efficient localizer to identify these regions for further evaluations.

We also aimed to identify a subregion selective to digits (ITG-numbers) in pITG by contrasting responses to digit against UFO stimuli, but found such a stimulus preference only in 35.30% and 11.76% of the participants in the left and right hemisphere, respectively. Given that here we used a narrow stimulus-set contrasting digits with only one other type of visual stimulus, more work is needed to determine if and how selectivity for digits can be found in pITG. This is particularly critical as some previous work successfully identifying ITG-numbers (Pinel et al. 1999; Roux et al. 2008; Cui et al. 2013; Shum et al. 2013; Hermes et al. 2015; Amalric and Dehaene 2016; Daitch et al. 2016; M. Grotheer et al. 2016; Mareike Grotheer et al. 2016) while other studies could not identify such a region (Pinel et al. 2004; Libertus et al. 2009; Cappelletti et al. 2010; Andres et al. 2012; Attout et al. 2014; Carreiras et al. 2015; Cummine et al. 2015; Holloway et al. 2015; Peters et al. 2015; Merkley et al. 2019) (for a meta-analysis see Yeo et al., 2017). Potential explanations for these differential results, include: 1) the number stimuli used (single digits, multiple digits, strings of digits etc.) and the stimuli numbers are contrasted with (faces, dot clouds, fourier noise, dice, etc.) vary drastically across experiments (Yeo et al. 2017), 2) the location of ITG-numbers is relatively close to the fMRI signal dropout zone of the ear canal, and specialized imaging acquisition parameters may be needed in this part of the brain (Shum et al. 2013; M. Grotheer et al. 2016), and 3) the task participants are performing vary drastically across studies and passive tasks in particular have been suggested to fail to elicit significant responses to digits in the pITG (Liu et al. 2025).

Interestingly, within ITG-math, we found differential sensitivities to stimulus features across hemispheres with larger overall responses to UFOs than digits in the univariate data as well as significant stimulus encoding in the multivariate analyses found only in the right hemisphere. These findings align with prior research showing subtle differences in the response profile of function regions in pITG across hemispheres. First, prior work found a positive correlation between age and neural responses for arithmetic only in the left but not in the right pITG (Rivera et al. 2005). Further, EEG research suggests a dynamical interplay between the bilateral pITG during numerosity tasks, where the right pITG attributes numerical meaning at early stages, then passes it on to the left hemisphere for information integration, and finally the operational momentum is supported by the bilateral pITG (Jang and Hyde 2020). Indeed, our observed stimulus sensitivity of the right ITG-math might reflect remnants of an initial right-lateralization for non-symbolic numerosities similar to the right IPS (for a meta-analysis see: Faye et al., 2019). However, others did not find such hemispheric asymmetries in pITG functional subregion (Hermes et al. 2015; Amalric and Dehaene 2016; Daitch et al. 2016; Grotheer et al. 2018) and further research investigating the division of labor of ITG-math across hemispheres is needed. There are several limitations of the current study that should be considered. First, sample size might have played a role in us not being able to identify ITG-numbers. However, it is important to note that here we aimed to identify the subregions in individual subjects and the relative proportion of identified ITG-numbers to ITG-math fROIs in this study, which was roughly 1:2 and 1:3 in the left and right hemispheres, respectively, is unlikely to change in a larger sample. Further, while we matched the numerosity and color tasks as closely as possible in terms of domain-general demands (working memory, attention, etc.) the observed differences in performance across tasks may suggest larger engagement in one task over the other. As such, more work is needed to confirm that the observed task preference reflects domain-specific numerosity processing. Our results are also limited to a numerosity task, which raises the question if ITG-math engages in other more complex mathematical tasks (e.g. addition, multiplication, subtraction and division). Prior work using arithmetical equations including addition and multiplication, as well as formulas (i.e. Zeta function) (Hermes et al. 2015; Amalric and Dehaene 2016; Amalric and Cantlon 2022) found increased activity in the pITG, thus suggesting that ITG-math may be engaged in various mathematical tasks. Similarly, here we used only two types of stimuli, digits and UFOs, yet numerosities can be represented by various other visual stimulus categories as well (Grotheer et al. 2018). Moreover, as we used only visual stimuli, our results are limited to the visual domain, yet prior work has suggested that pITG is involved in mathematical processing even in other modalities (Abboud et al. 2015; Amalric and Dehaene 2016). As such, future work employing a larger variety of stimuli including (e.g. number words or Roman numerals), different sensory modalities and experimental paradigms is needed to fully grasp the functional role of pITG subregions.

We conclude that the bilateral ITG-math, which can be consistently found across individuals, is involved in numerosity processing independent of the visual features of the presented stimulus. Moreover, we propose that the developed experimental paradigm, which we made openly available, may serve as a localizer to identify numerosity-selective responses in the pITG and to hence facilitate further research on these regions.

## Acknowledgements

Funded by the European Union (ERC, project WRAPPED, 101161197). Views and opinions expressed are however those of the author(s) only and do not necessarily reflect those of the European Union or the European Research Council. Neither the European Union nor the granting authority can be held responsible for them. This work was further supported by the Deutsche Forschungsgemeinschaft (German Research Foundation, DFG) under Germany’s Excellence Strategy (EXC 3066/1 “The Adaptive Mind”, Project No. 533717223) and by the Deutsche Forschungsgemeinschaft (DFG, German Research Foundation) - project number 222641018 - SFB/TRR 135 TP C10, as well as by a LOEWE Professorship awarded by the State of Hesse (LOEWE/4b//519/05/01.002(0016)/120). MR-imaging for this study was performed at the Bender Ins⊠tute of Neuroimaging (BION) at the Justus Liebig University Giessen, Germany.

