## Supplementary figures for "Regions in the human inferior temporal gyrus are engaged in numerosity processing across visual stimulus categories"

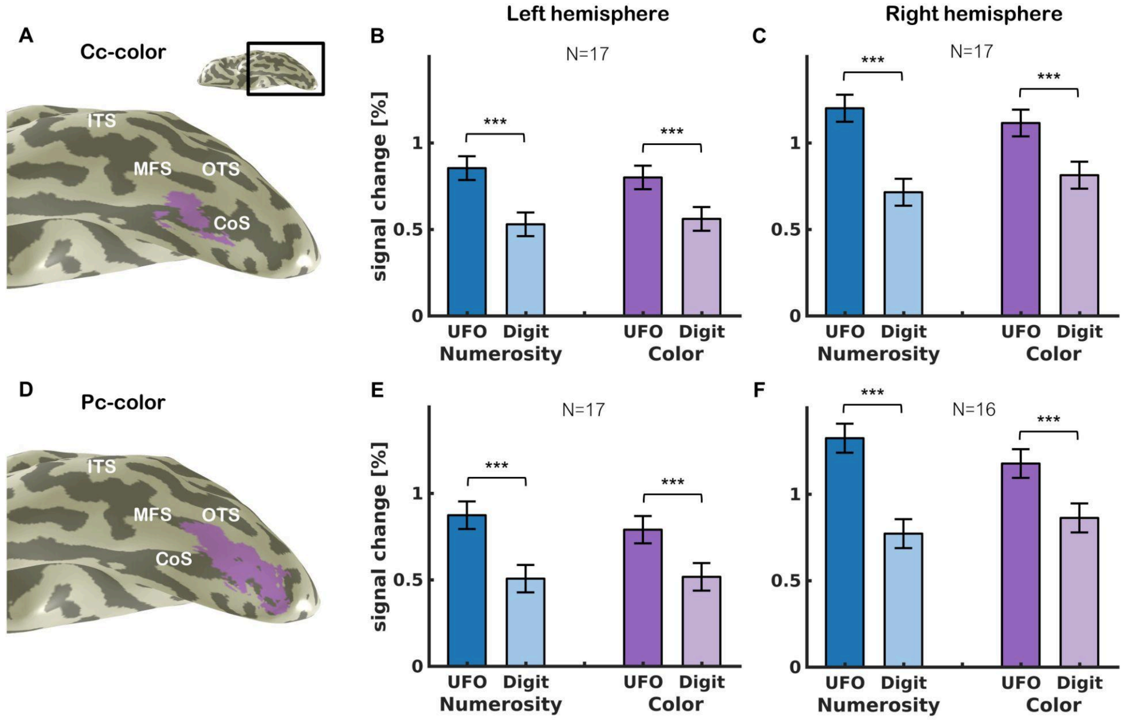


**Supplementary Fig. 1. Bilateral Cc- and Pc-color regions show a preference for UFO stimuli and no task preference.** A. Cc-color: voxels in the median fusiform gyrus (FG) and collateral sulcus (CoS) that showed significantly higher responses for colored than gray-scale images in the color localizer (T=3, voxel level, uncorrected). Data is shown in a ventral view of the inflated left hemispheres of a representative subject. B. Mean responses ± SEM across 17 subjects in the left Cc-color from independent data. C. Mean responses ± SEM across 17 subjects in the right Cc-color from independent data. D. Pc-color: voxels in the median fusiform gyrus (FG) and collateral sulcus (CoS) that showed significantly higher responses for colored than gray-scale images in the color localizer (T=3, voxel level, uncorrected). E. Mean responses ± SEM across 17 subjects in the left Pc-color from independent data. F. Mean responses ± SEM across 16 subjects in the right Pc-color from independent data. Asterisks indicate main effect of stimulus across hemispheres, ***P<0.001. Abbreviations: ITS: inferior temporal sulcus, OTS: occipitotemporal sulcus, MFS: midfusiform sulcus, CoS: collateral sulcus.

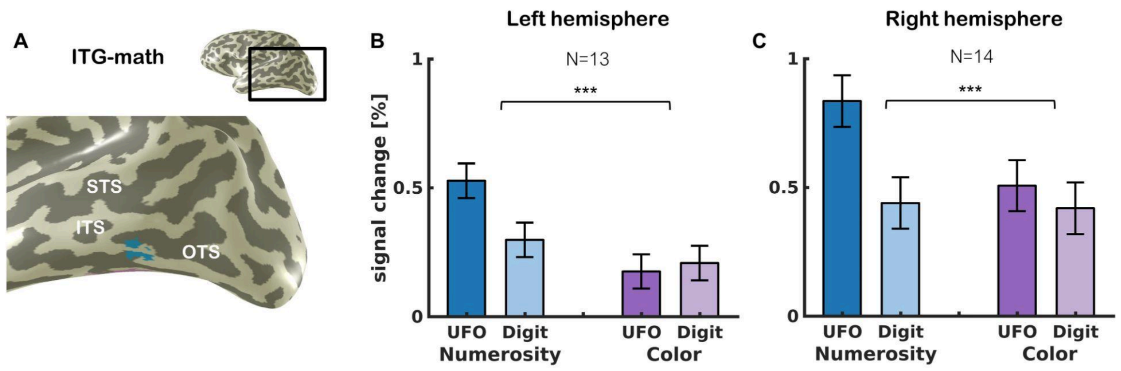


**Supplementary Fig. 2. ITG-math defined from data from the first two runs shows a preference for the numerosity task.** A. ITG-math: voxels in the ITG/ITS that showed significantly higher responses during the numerosity than the color task (T=3, voxel level, uncorrected). Data is shown in the lateral view of the inflated left hemisphere of a representative subject. B. Mean responses ± SEM across 13 subjects in the left ITG-math from independent data. C. Mean responses ± SEM across 14 subjects in the right ITG-math from independent data. Asterisks indicate main effect of task across hemispheres, ***p<0.001. Abbreviations: STS: superior temporal sulcus, ITS: inferior temporal sulcus, OTS: occipitotemporal sulcus.

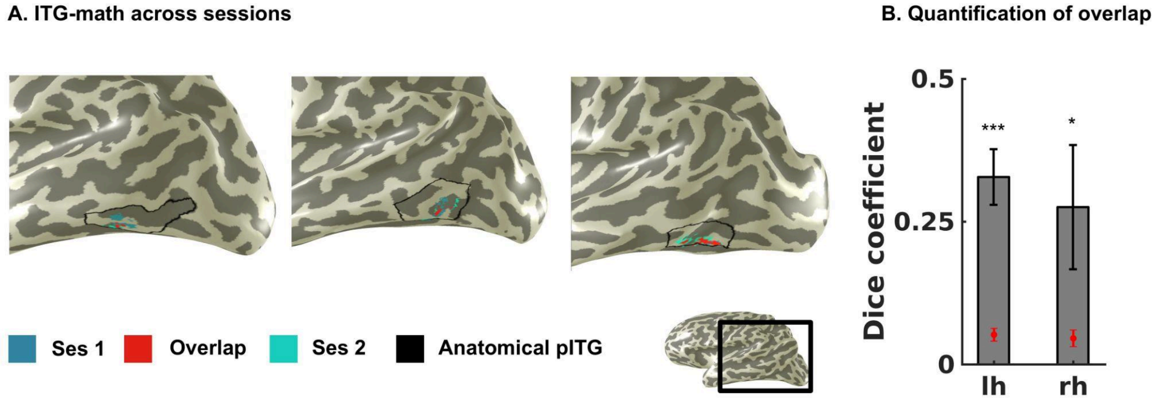


**Supplementary Fig. 3. ITG-math fROIs based on data from two runs show consistent localization across sessions.** A. Figure shows the inflated cortical surface of three participants representing examples of low (left), medial (middle), and high (right) spatial overlap in ITG-math fROIs across sessions. B. Quantification of the overlap between fROIs across sessions using the dice coefficient (DC), mean across subjects ± SEM. Red dots indicate chance level. Star indicates DCs is significantly higher than chance, *p<0.05, ***p<0.001.

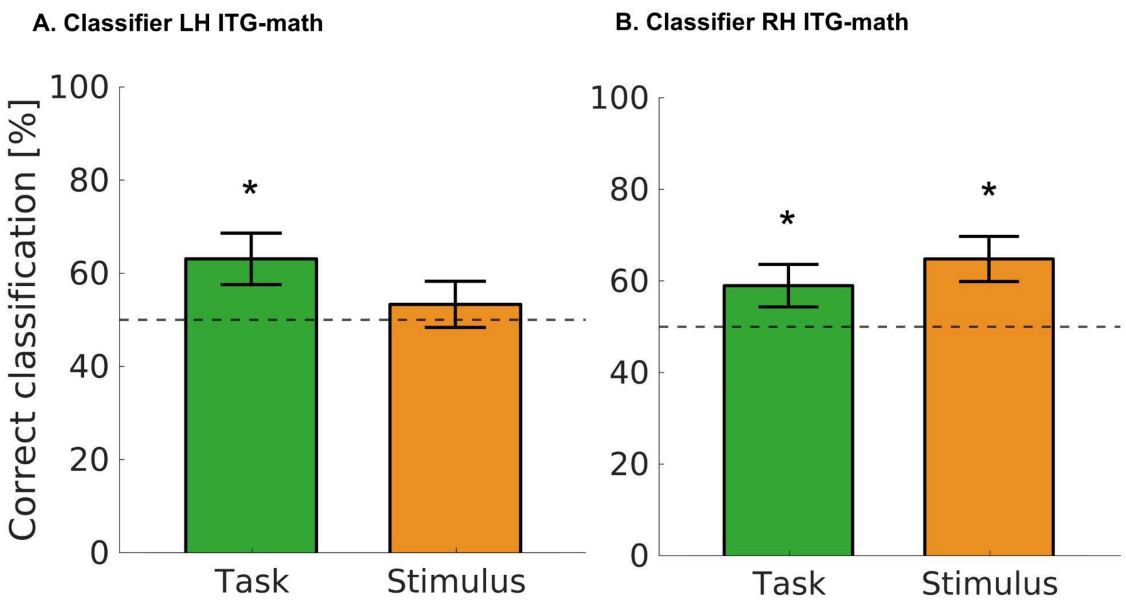


**Supplementary Fig. 4. Decoding accuracies in ITG-math varies across conditions and hemispheres.** Mean ± SEM winner-takes-all (WTA) classification performance for task and stimulus in left (A) and right (B) ITG-math. Star indicates that classifier performance is significantly above chance level (p<0.01 with Bonferroni-corrected threshold). Abbreviations: RSM, representational similarity matrix; WTA, winner-takes-all.

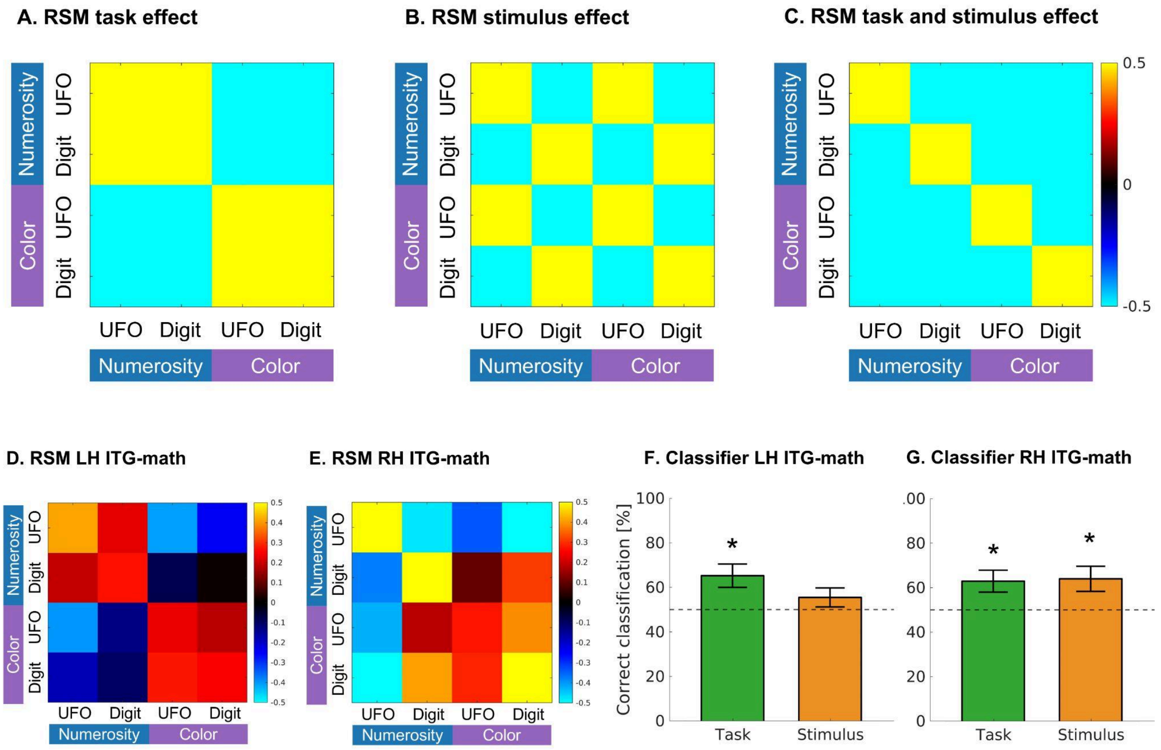


**Supplementary Fig. 5. Distributed responses in native ITG-math fROIs vary across hemispheres.** A-C.

Model RSMs depicting hypothetical similarities across conditions based on task (A), stimulus (B), and combined task and stimulus effects (C). **D-E**. RSM from ITG-math in the left (D. N=16) and right (E. N=15) hemispheres. Conditions are arranged by stimulus (UFO vs. digit) and grouped by task (numerosity vs.color). In the left hemisphere, significant correlations with task (r=0.87, p<0.0001), and combined task and stimulus models (r=0.66, p=0.006) were observed, but not with stimulus model (r=0.11, p=0.69). In the right hemisphere, stimulus (r=0.54, p=0.03) and combined task and stimulus models (r=0.63, p=0.009) showed significant correlations, but not task model (r=0.43, p=0.09). F-G. Mean ± SEM winner-takes-all (WTA) classification performance for task and stimulus in left (F) and right (G) ITG-math. Star indicates that classifier performance is significantly above chance level (p<0.01 with Bonferroni-corrected threshold). Abbreviations: RSM, representational similarity matrix; WTA, winner-takes-all.

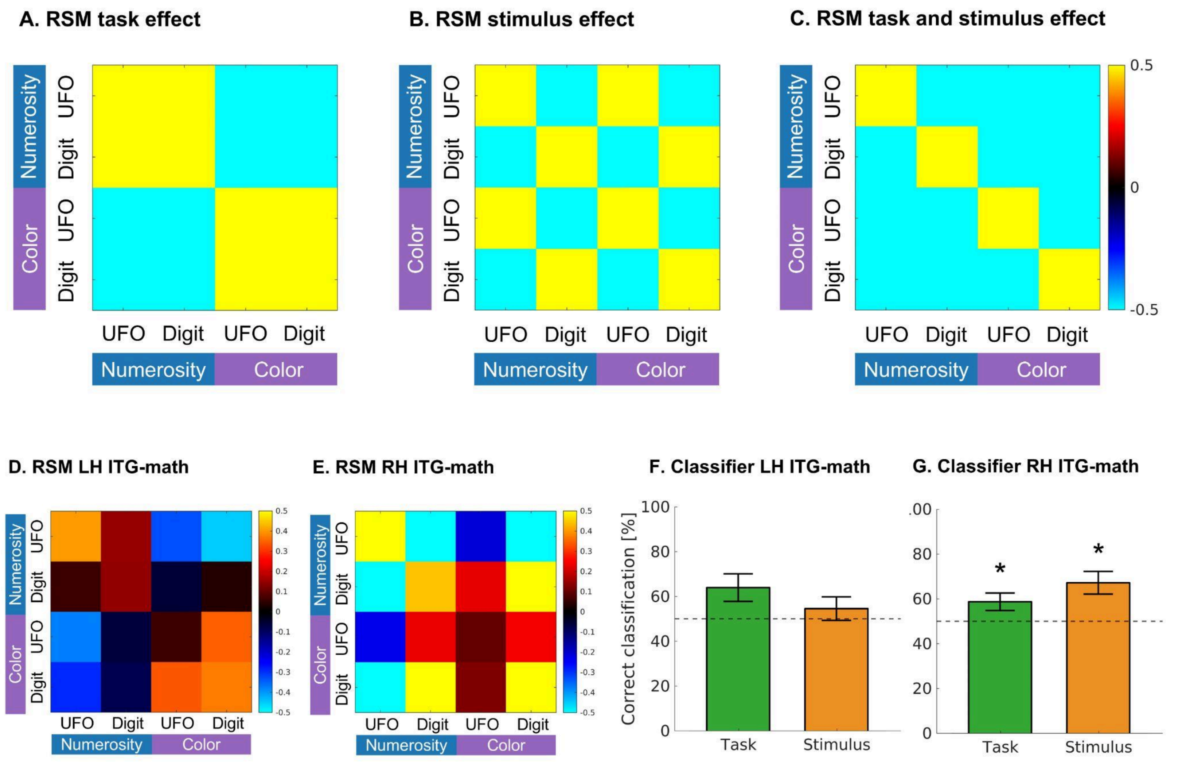


**Supplementary Fig. 6. Distributed responses in 2-run ITG-math vary across hemispheres.** A-C. Model RSMs depicting hypothetical similarities across conditions based on task (A), stimulus (B), and combined task and stimulus effects (C). **D-E**. RSM from ITG-math in the left (D. N=13) and right (E. N=14) hemispheres. Conditions are arranged by stimulus (UFO vs. digit) and grouped by task (numerosity vs.color). In the left hemisphere, significant correlations with task (r=0.87, p<0.0001), and combined task and stimulus models (r=0.56, p=0.02) were observed, but not with stimulus effect (r=0.05, p=0.84). In the right hemisphere, only stimulus model (r=0.54, p=0.03) showed significant correlations (task: r=0.11, p=0.69; combined task and stimulus: r=0.44, p=0.09). F-G. Mean ± SEM winner-takes-all (WTA) classification performance for task and stimulus in left (F) and right (G) ITG-math. Star indicates that classifier performance is significantly above chance level (p<0.01 with Bonferroni-corrected threshold). Abbreviations: RSM, representational similarity matrix; WTA, winner-takes-all.

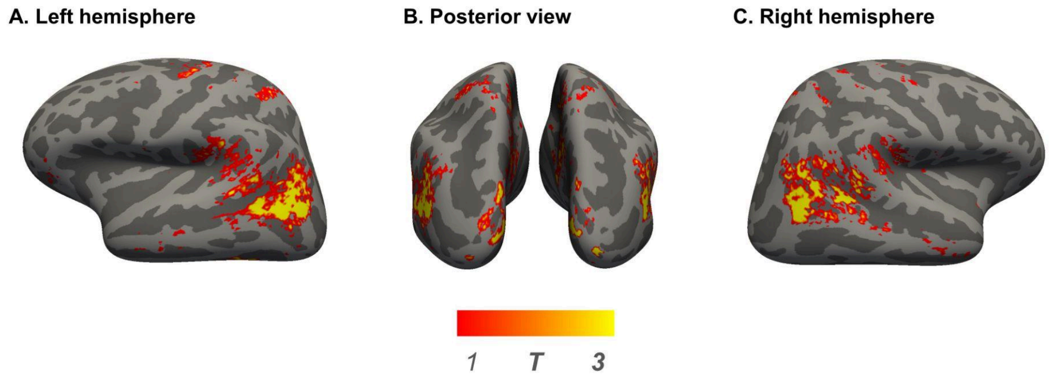


**Supplementary Fig. 7. Whole-brain analysis contrasting digit and UFO stimuli across participants.** Responses elicited by contrasting the digit with the UFO stimuli in both sessions were averaged across participants and are presented on the inflated cortical surface of the Freesurfer average brain.

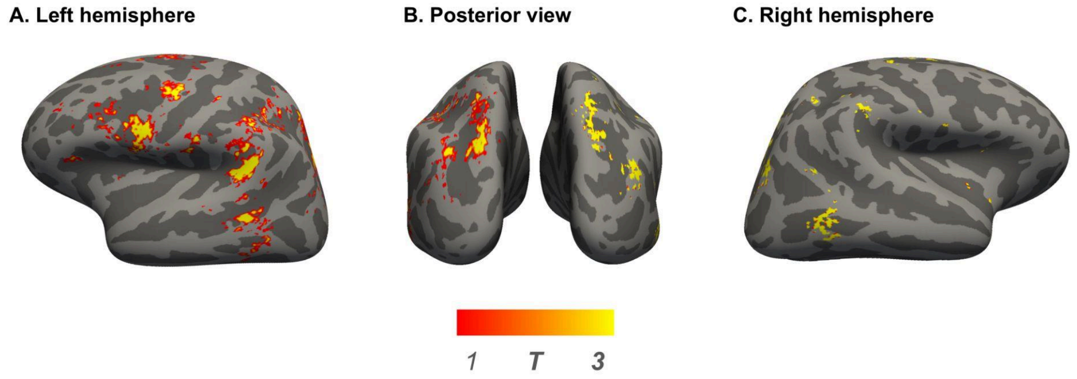


**Supplementary Fig. 8. Whole-brain analysis contrasting the numerosity and color tasks across participants using only 2 runs of data.** Responses elicited by contrasting the numerosity with the color task in two runs of session 1 were averaged across participants and are presented on the inflated cortical surface of the Freesurfer average brain.

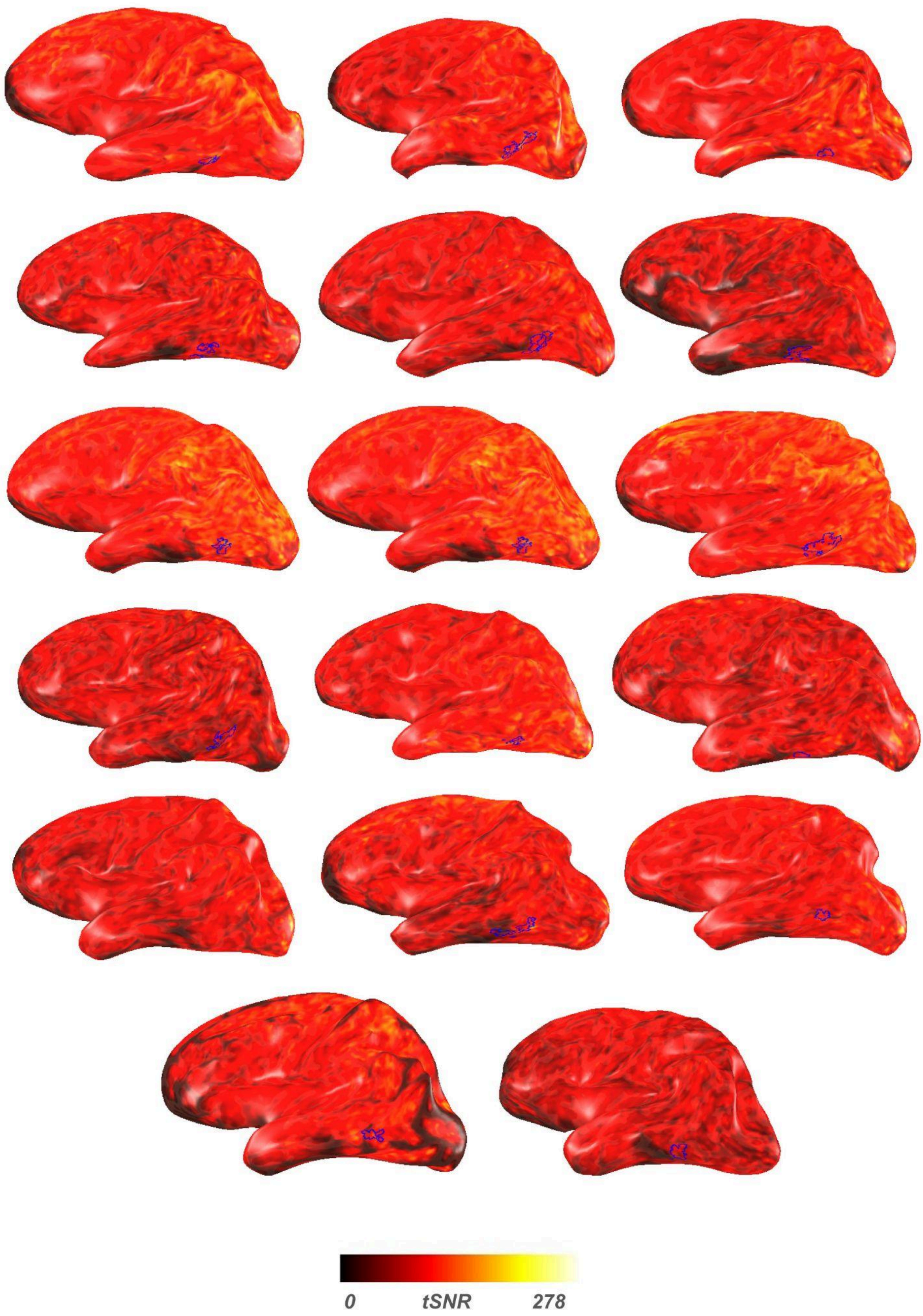


**Supplementary Fig. 9. Left ITG-math lies outside the signal dropout zone.** Inflated left hemispheres of all participants in native brain space showing the average temporal signal-to-noise ratio (tSNR) from Ses 1. The outline of ITG-math (blue) is superimposed where it could be defined. As can be observed, ITG-math is adjacent to, but not within, the signal dropout zone near the petrous bone.

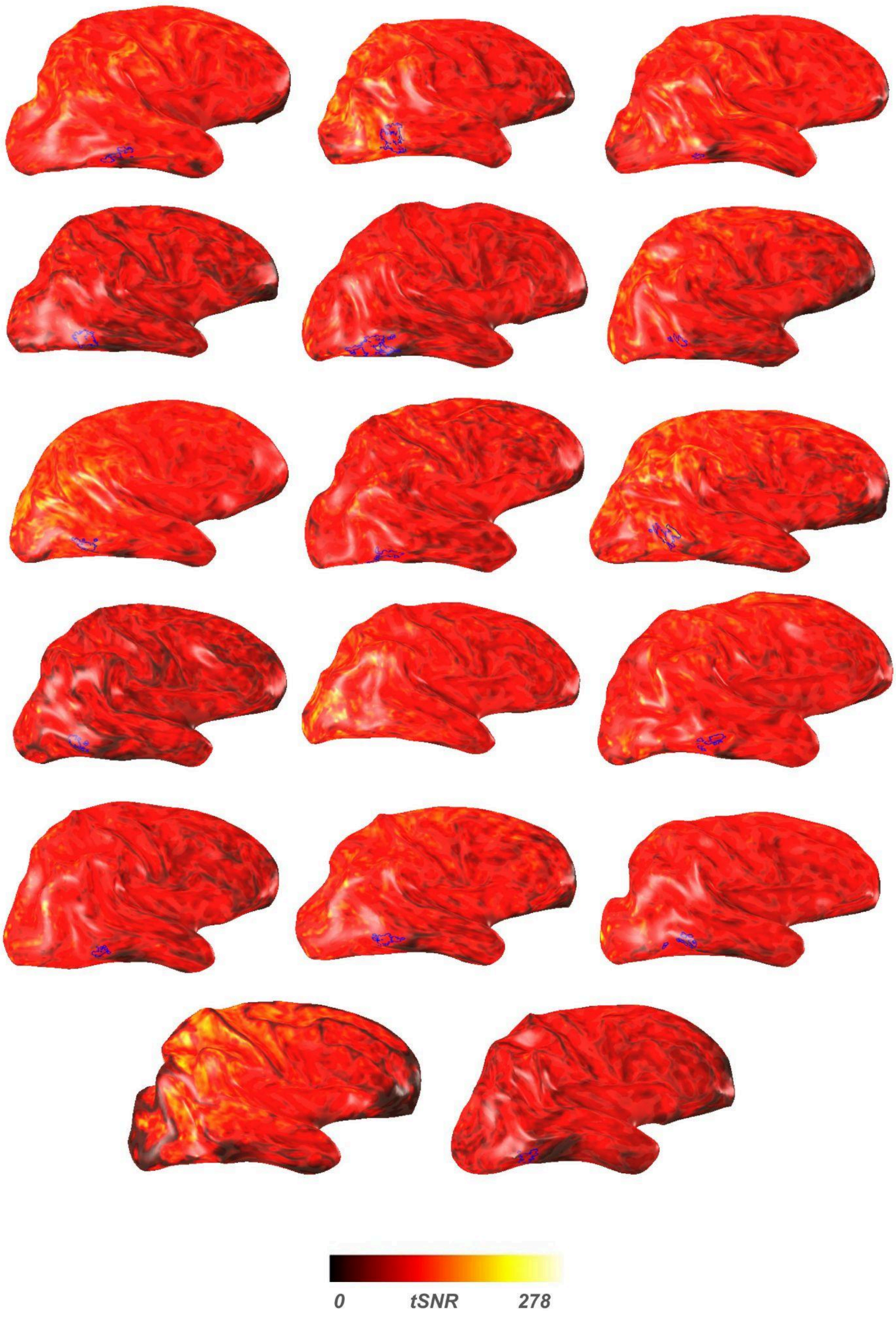


**Supplementary Fig. 10. Right ITG-math lies outside the signal dropout zone.** Inflated right hemispheres of all participants in native brain space showing the average temporal signal-to-noise ratio (tSNR) from Ses 1. The outline of ITG-math (blue) is superimposed where it could be defined. As can be observed,

ITG-math is adjacent to, but not within, the signal dropout zone near the petrous bone.
